# The *Pseudomonas putida* Type VI Secretion Systems Shape the Tomato Rhizosphere Microbiota

**DOI:** 10.1101/2025.08.14.670259

**Authors:** David Vázquez, Cristina Civantos, David Durán-Wendt, Adrián Ruiz, Rafael Rivilla, Marta Martín, Patricia Bernal

## Abstract

Bacterial competition mechanisms drive microbial community dynamics across diverse ecological niches. The Type VI Secretion System (T6SS) represents a sophisticated nanomachine used by Gram-negative bacteria for contact-dependent elimination of competitors through the delivery of toxic effectors. While the T6SS has been well-documented in mammalian gut microbiota development, its role in shaping plant rhizosphere communities remains poorly understood despite the ecological importance of rhizosphere microbiota. This study investigates how the three *Pseudomonas putida* KT2440 T6SS clusters influence the tomato rhizosphere microbiota in agricultural soil. Through comprehensive *in vitro* and *in vivo* analyses, we demonstrate that while the K2/K3-T6SSs remain inactive under standard laboratory conditions, they become specifically activated in the presence of plant pathogens, suggesting an adaptive response to competitive pressure. Our experiments with T6SS-deficient mutants reveal that the *P. putida* T6SSs are essential for effective rhizosphere colonisation, with mutant strains showing significantly reduced colonisation capabilities compared to wildtype strain in competitive soil environments. Most importantly, our data establish that the *P. putida* T6SSs directly shape the taxonomic diversity and community structure of the rhizosphere microbiota of tomato plants. These results place the T6SS as a critical factor driving the evolution of complex polymicrobial communities within the plant rhizosphere, paralleling its established role in the gut microbiota. This research advances our understanding of the ecological functions of the different T6SSs in *P. putida* and the molecular mechanisms underlying microbial community assembly in the rhizosphere. Thus, it offers valuable insights for agricultural applications involving beneficial microbes and plant health management strategies.

## INTRODUCTION

Bacteria commonly coexist with other microorganisms, forming intricate communities characterised by dynamic interactions that include cooperation and competition. To survive in these harsh and diverse environmental niches, they have developed different competition mechanisms to combat foes and manipulate host cells [1]. The polymicrobial communities associated with relevant hosts, such as plants, insects, and mammals, that is, the microbiota, are especially dense and complex, and competition mechanisms are frequent among their members. Over the past decades, knowledge regarding the weaponry arsenal used for bacterial competition has expanded dramatically, ranging from direct release of effectors to intricate nanomachines with fascinating structures and mechanisms of action [2]. Among these sophisticated nanomachines is the type VI secretion system (T6SS), a multiprotein contractile apparatus employed by gram-negative bacteria. The T6SS is used to eliminate competitors through contact-dependent secretion of a diverse range of toxins. Most T6SS effectors delivered by this lethal machine are designed to kill rivals, but to a lesser extent, they can be responsible for exerting social control, subverting host cells, and controlling access to common goods, such as key nutrients [3]. Thus, the T6SS is considered an influential mechanism driving the evolution of complex polymicrobial communities, such as the gut and rhizosphere microbiota [4].

Over the last decades, numerous studies have revealed the significant impact of the microbiota on the health of its host, especially the mammalian gut microbiota. The mammalian gastrointestinal tract harbours a very dense microbial community dominated by four bacterial phyla, *Actinobacteria*, *Bacteroidetes*, *Firmicutes*, and *Proteobacteria* [5], where the T6SSs are widely distributed [6].

The T6SS also influences the insect gut microbiota, for example, *Pieris brassicae* is a cabbage pest that can be suppressed by the biocontrol agent *Pseudomonas protegens* CHA0. *P. protegens* CHA0 uses the T6SS to outcompete members of the insect microbiota, mainly members of the *Enterobacteriaceae* family and thus promotes colonisation and lethal infection in the host, protecting cabbage crops [7]. Additionally, the T6SS plays an important role in the evolution of the gut microbiota of honey and bumble bees through diverse T6SS effectors, with the potential for inter-strain transfer among bee gut bacteria [8, 9]. The T6SS is considered a crucial factor in maintaining the equilibrium between the mechanisms of host protection via pathogen exclusion and the risk of host infection due to the elimination of commensal species. Currently, there is a solid consensus in the field on the essential role of the T6SS in shaping the gut communities. However, our understanding of its involvement in other key ecosystems, such as the rhizosphere, remains limited.

The gut and rhizosphere are both eukaryotic systems hosting complex polymicrobial communities [10]. Therefore, it is plausible to suggest that the plant root microbiota is also modulated by the T6SS. Many plant-related bacteria, such as phytopathogens, *e.g. Agrobacterium tumefaciens, Pseudomonas syringae*, *Pseudomonas savastanoi* and *Pectobacterium atrosepticum*, or the biocontrol agents *Pseudomonas putida*, *Pseudomonas fluorescens* MFE01 and *Pseudomonas ogarae* F113 contain T6SSs [11, 12]. The T6SSs of bacterial phytopathogens may not directly enhance virulence towards plant cells, but they facilitate microbial competition, leading to improved fitness and persistence within the plant microbiota. This ultimately increases the probability that these bacteria cause diseases in their host plants [13–15] and only a few cases have been related to host manipulation [15, 16]. On the other hand, well-established commensal or biocontrol non-pathogenic bacteria mostly from the *Pseudomonas* genus, such as *P. putida*, *P. protegens, P. fluorescens* and *P. ogarae* successfully use the T6SSs to outcompete and inhibit plant pathogens and other putative rivals in the rhizosphere [7, 11, 12, 17–22]. Besides, in some rhizobia species, the T6SS play a role in the symbiosis process with the leguminous plant [23, 24]. These systems are active in environments or condition-mimicking environments that are ecologically relevant for these strains, that is, plant-like conditions for *A. tumefaciens, Sinorhizobium fredii, Azospirillum brasilense,* and *P. atrosepticum*, or they are constitutively active, such as the *P. putida* K1-T6SS [21, 24–27].

Despite our understanding of the T6SSs in individual plant-associated bacteria, the influence of this deadly system on the rhizosphere microbiota remains poorly understood. Initial investigations in *P. fluorescens* and *Kosakonia sp.* revealed that T6SS mutants exhibit a significantly reduced capacity to colonise both the rhizosphere and the endosphere [12, 28], and the opposite effect has been shown on the colonisation of watermelon cotyledons by *Acidovorax citrulli* [29]. These findings suggest a potential role of the T6SSs in shaping these microbial communities, as evidenced by a metagenomic analysis of an endophytic bacterial community inoculated with the plant pathogen *Burkholderia glumae*. The study showed significant differences in the overall taxonomic diversity between the microbiota inoculated with the wildtype and that inoculated with the T6SS mutant [30].

To gain a better understanding of the role of the T6SSs in natural environments, such as the rhizosphere of plants, the present study comprehensively examined the contribution of the *P. putida* T6SSs to the native microbiota within the rhizosphere of tomato plants grown in agricultural soil. *P. putida* is an efficient biocontrol agent [31, 32] that contains three T6SSs, but only one has been previously detected to be active under laboratory conditions [17, 27]. The K1-T6SS is used to eliminate ecologically relevant competitors, including severe plant pathogens such as *P. syringae*, *P. savastanoi* and *X. campestris* [17, 18]. These pathogens are the leading causes of deadly diseases in many economically important crops, such as carrots, potatoes, and olive trees [33], illustrating the importance of biocontrolling them. In this study, we demonstrate for the first time the functionality of the K2 and/or K3-T6SS clusters *in vitro* in the presence of a plant pathogen and *in vivo* in the presence of the rhizosphere microbiota. Here, we have also shown the importance of the T6SS in *P. putida* KT2440 rhizosphere competitive colonisation. These data indicate the importance of the T6SSs in the evolution of a natural, complex, and mixed polymicrobial community, such as that present in the rhizosphere of tomato plants.

## MATERIALS AND METHODS

### Reagents and bacterial growth conditions

Unless otherwise stated, chemicals, antibiotics and reagents were acquired from Sigma Aldrich. Lysogeny broth Lennox (LB) (5 g/L NaCl) and agar (1.5% w/v) [34] were used for routine growth of all organisms with shaking at 200 RPM, as appropriate; *Escherichia coli* was grown at 37 °C, *P. putida* at 30 °C and *Xanthomonas campestris* at 28 °C. SA medium (20 g/L sucrose, 2 g/L _L_-asparagine, 1.31 g/L K_2_HPO_4_·3 H2O, 0.5 g/L MgSO_4_·7 H20 and 15 g/L agar; pH 7.0) [35] was used to recover bacteria from soil. Vogel-Bonner Medium (VBM) (200 mg/L MgSO_4_.7H_2_O; 2 g/L anhydrous citric acid; 10 g/L K_2_HPO_4_; 3.5 g/L NaNH_4_HPO_4_.4H_2_O; pH 7.0) was used to select transconjugants after triparental conjugation, as *E. coli* donor and helper cells cannot grow using citrate as carbon source. Growth media were supplemented with the following antibiotics, as required: 20 μg/mL rifampicin for *P. putida*; 20 μg/mL gentamicin, for *E. coli* and *X. campestris*; 25 μg/mL kanamycin for *E. coli;* 400 μg/mL streptomycin *P. putida* and 50 μg/mL for *E. coli* and 30 μg/mL chloramphenicol for *E. coli*.

### Construction of bacterial strains

The strains, plasmids and primers used in this study are depicted in supplementary Tables S1, S2 and S3, respectively. *P. putida* gene mutants (*tssM2 and tssM2 tssM3*) were constructed by allelic exchange, as previously described [17]. Briefly, 500-bp DNA fragments upstream and downstream of the gene to be deleted were amplified using *P. putida* KT2440 genomic DNA. A fragment containing both regions was obtained by overlapping PCR, cloned into pCR-BluntII-TOPO (Invitrogen), and the DNA was sequenced and subcloned into the *Xba*I/*Bam*HI sites of pKNG101 [17]. We used primers P1-P6 for *tssM2* mutation and P7-P12 for *tssM3.* Primers P13-P14 and P15-P16 were used to sequence fragments inserted into pTOPO and pKNG101, respectively (see supplementary Table S3). The suicide vector pKNG101 [36] does not replicate in *Pseudomonas*; it was maintained in *E. coli* CC118λpir and mobilised into *Pseudomonas* by triparental conjugation [37]. The mutants in which double recombination events occurred, resulting in the deletion of the gene of interest, were selected on sucrose plates as previously described [36]. Finally, the deletion event was confirmed by PCR using the primers described in Table S3 and subsequently sequenced by Nanopore Technology (Plasmidsaurus) to confirm there were no additional mutations.

### Interbacterial competition assays

*In vitro* competition assays were performed on LB (5 g/L NaCl) agar (1.5% w/v) plates, as previously described [38]. Briefly, overnight bacterial cultures were washed and adjusted to an OD_600_ of 10 in sterile PBS and mixed in a 1:1 ratio (*P. putida*:prey). Mixtures were grown on LB agar plates at 30°C for 5 hours (*E. coli* prey) or 24 hours (plant pathogen as prey) and then collected using an inoculating loop and resuspended in sterile PBS. The outcome of the competition was quantified by counting colony forming units (CFUs) using antibiotic selection of the input (time = 0 hours) and output (time = 5 hours or time = 24 hours). All prey strains harboured the plasmid pRL662, which confers resistance to gentamicin and was used for antibiotic selection. The *P. putida* KT2440 strain used in this study is naturally resistant to rifampicin (KT2440R), and thus, this antibiotic was used for *P. putida* selection. For all competition assays, at least three biologically independent experiments were performed. Competitive index values were calculated using the following formula:

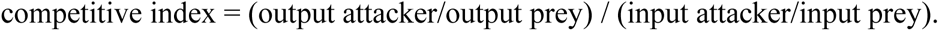

### Growth curve assay

To perform the growth curves, overnight cultures of each strain were grown in LB supplemented with the appropriate antibiotic. The next day, the cultures were adjusted to OD_600_ 0.05 in the same medium. Aliquots of 200 μL were placed in a microtiter plate, and the Synergy/H1 microplate reader (Biotek©) was used to measure the OD_600_ every 15 min at 30 °C for 10 hours. Data was obtained for three biological replicates, each with technical duplicates.

### Rhizosphere colonisation analysis

To evaluate rhizosphere colonisation of tomato plants (Rebelion F1, Vilmorin, France) by the wildtype strain *P. putida* KT2440 and its T6SS mutant derivates, seeds underwent a sterilisation process. First, the seeds were immersed in 70 % ethanol for 1 min, followed by 5 % sodium hypochlorite immersion for a further 3 min and subsequent washing with sterile distilled water. After thorough washing, the seeds were germinated on sterile 1 % agar plates maintained at 28 °C for 24 hours. The germinated seedlings were then individually transferred to sterile 50 mL polypropylene tubes (FALCON®). Each tube contained 16 g of a 1:1 mixture of non-sterile agricultural soil (Aranjuez, Spain – GPS coordinates 40.025, −3.684) and sterile sand (Merck KGaA, Darmstadt, Germany), supplemented with 3 mL of sterile water. The tubes were incubated in a controlled environment room set at 25 °C with a 16-hour light cycle to promote growth for 7 days. After this, each seedling was inoculated with 1 mL of a bacterial culture (10^6 CFU/mL) consisting of the wildtype *P. putida* KT2440, and a single (*tssA1*), a double (*tssM2 tssM3*) or a triple mutant strain (*tssA1 tssM2 tssM3*). The plants were then cultivated for an additional two weeks. Post incubation, the aerial parts of the plants were removed, and 10 mL of a 0.85 % sterile saline solution was added to each tube. The tubes were vortexed thoroughly to resuspend the bacteria present in the rhizosphere. The resulting suspension was decanted to separate the soil and sand matrix, and serial dilutions of the supernatant were plated on SA plates supplemented with rifampicin (100 μg/mL). Each condition was tested in six replicates, with non-inoculated plants as negative controls.

### DNA Extraction & Sequencing

Soil DNA was extracted using the FastDNA Spin Kit for Soil (MP Biomedicals, USA), following the manufacturer’s protocol with additional modifications [39]. For bacterial 16S rRNA profiling, the extracted soil DNA samples were sent to Macrogen. The V3-V4 regions of the 16S rRNA gene were amplified using Illumina MiSeq paired-end 2×300 system with primers P17 (341F) and P18 (805R) (Table S3) [40]. A rarefaction curve was employed to assess whether the sequencing depth was sufficient to capture the majority of the bacterial taxa present in the samples. As observed in Fig. S1, all curves plateaued, indicating that sequencing saturation was reached, and the observed diversity approached the true diversity for all analysed samples.

### ASVs determination and bacterial diversity analyses

Demultiplexed files containing raw reads were processed using the DADA2 Pipeline (https://benjjneb.github.io/dada2/tutorial.html). The reads underwent quality filtering, denoising, clustering into amplicon sequence variants (ASVs), and chimaera removal using the DADA2 v1.18 package in R [41]. The ASV data, the nucleotide sequences and the abundance were exported to QIIME2 v2023.2.0 [42] for further analysis. In QIIME2, the ASVs were aligned using MAFFT [43, 44] and a phylogenetic tree was constructed with FastTree2 [45]. Taxonomic assignment was performed using the q2-feature-classifier [46] based on the SILVA 99% 16S sequence database, release 138 [47]. The SILVA database sequences were trimmed to match the amplicons used in this study. A naive Bayes classifier [48] was constructed from the processed database, and the taxonomic assignment of ASVs was carried out using the QIIME2 classify-sklearn tool [49].

Alpha-diversity (within-sample diversity) and beta-diversity (between-sample diversity) analyses were conducted using the QIIME2 diversity plugin. Alpha-diversity was assessed for evenness and richness using the Shannon Index. The significance of the replicate effect on alpha-diversity was evaluated using the Kruskal-Wallis test. Beta-diversity was estimated using Bray-Curtis distances [50] between samples. To identify differential populations between samples, the ANCOM-BC package [51] was employed, selecting those ASVs with a log_2_ fold change greater than 1,5 and an adjusted p-value of ≤ 0.001. The ANCOM-BC package excludes all taxa below 0.1% abundance.

### Statistical analysis of experimental data

Statistical analysis was performed in GraphPad PRISM v8.0.2 using an unpaired two-tailed T-test. Statistical significance was defined as p < 0.05. Statistical analysis for the quantification of bacterial recovery from rhizosphere colonisation was performed in R v4.3.2. The normal distribution of data was checked with the Shapiro-Wilk test. Data were analysed with the Kruskal-Wallis non-parametric test due to non-normal distribution. For diversity analysis, data were checked with the PERMANOVA test in QIIME2.

## RESULTS

### The K2/K3-T6SSs are not used to kill standard laboratory strains

*P. putida* KT2440 is equipped with three T6SSs named K1-, K2- and K3-T6SSs. The K2- and K3-T6SSs are phylogenetically related, both belonging to the T6SS phylogenetic group 1.2, while the K1 cluster clades separately and it belongs to T6SS phylogenetic group 4B [11, 17]. The K1-T6SS system is active under laboratory conditions and has been proven to kill *E. coli* and resilient plant pathogens [17, 18]. The K2- and K3-T6SSs contain 12 of the 13 genes encoding core T6SS components, missing *clpV*, the gene that encodes the ATPase required for the disassembling of the sheath (Fig. 1).

**Figure 1.**
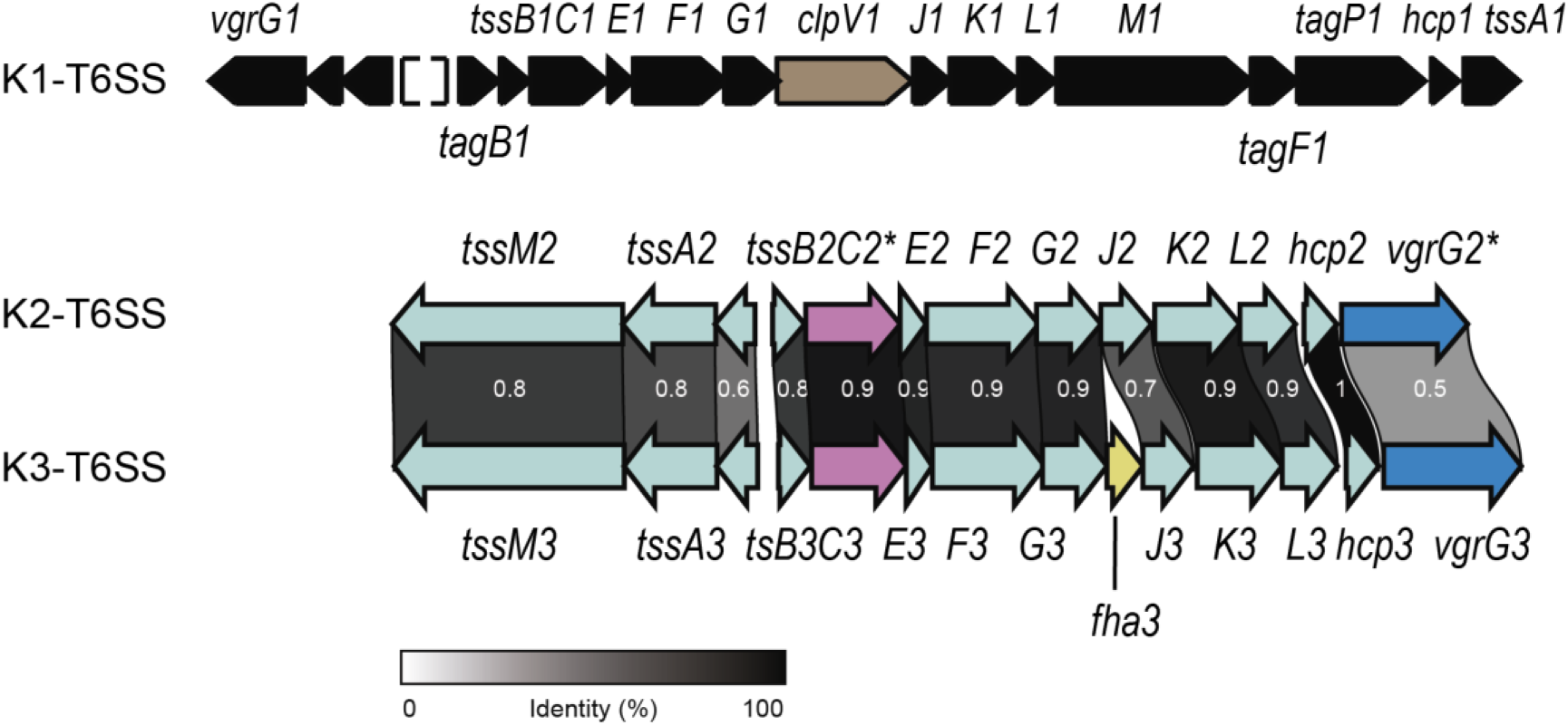
The *P. putida* T6SSs genetic clusters. Schematic representation of the genomic organisation of the structural genes of the K1-, K2- and K3-T6SS clusters from *P. putida*. For clarity, the clusters do not include the genes found downstream of the *vgrG* genes. K1-T6SS genes are shown in black, except *clpV1,* which is illustrated in brown. K2- and K3-T6SSs genes are depicted in light green except for pseudogene *tssC2\** and gene *tssC3* shown in pink and pseudogene *vrgG2\** and gene *vgrG3* in blue. The gene encoding the regulatory element *fha3*, present exclusively in the K3-T6SS cluster, is coloured in yellow.

The K2-T6SS cluster contains two pseudogenes, *vgrG2\** and *tssC2\**, which present premature stop codons (Fig. 1, in pink and blue respectively) [17], implying that this system is not functional. However, the genome of *P. putida* KT2440 contains several genes encoding related VgrGs, *i.e.* VgrG3, VgrG4 and VgrG5, and TssC proteins, *i.e.* TssC3, that could be shared between different T6SSs. This could be expected because the amino acid identity of these proteins is remarkably high, ranging from 71% to 96% (Fig. S2), as is the rest of the proteins of these two clusters [17]. Among the core components essential for T6SS activity, TssM is involved in the first stages of system assembly as a membrane complex component and TssA is the orchestrator of baseplate localisation onto the membrane complex and primes tail polymerisation [3]. Thus, mutants in either of the genes that encode TssM or TssA do not have functional T6SSs. Since the functionality of the K2 system is doubtful, we constructed a double mutant *tssM2 tssM3* to disable *P. putida* K2- and K3-T6SSs, along with a *tssA1* mutant to inactivate the K1-T6SS [17]. Additionally, we used a triple mutant *tssA1 tssM2 tssM3* with the three systems knocked out [17]. We performed *in vitro* competition assays using *E. coli* K12 as prey and *P. putida* wildtype or T6SS mutants as predators to reveal whether these clusters are active under these laboratory conditions. As previously reported, the competitive index of the single *tssA1* mutant was reduced compared to the wildtype strain [17, 18]. We observed the same effect in the triple mutant, but not in the *tssM2 tssM3* double mutant (Fig. 2). These data suggest that only the absence of *tssA1* is responsible for the reduction in the killing capabilities of these strains. Moreover, we found that compared to a wildtype strain, the double mutant *tssM2 tssM3* exhibited a similar competitive index when cocultured with *E. coli* (Fig. 2). Since the phenotype of the double K2/K3-T6SS mutant was identical to the wildtype strain, we conclude that the K2- and K3-clusters are not used to kill *E. coli* under these experimental conditions (Fig. 2).

**Figure 2.**
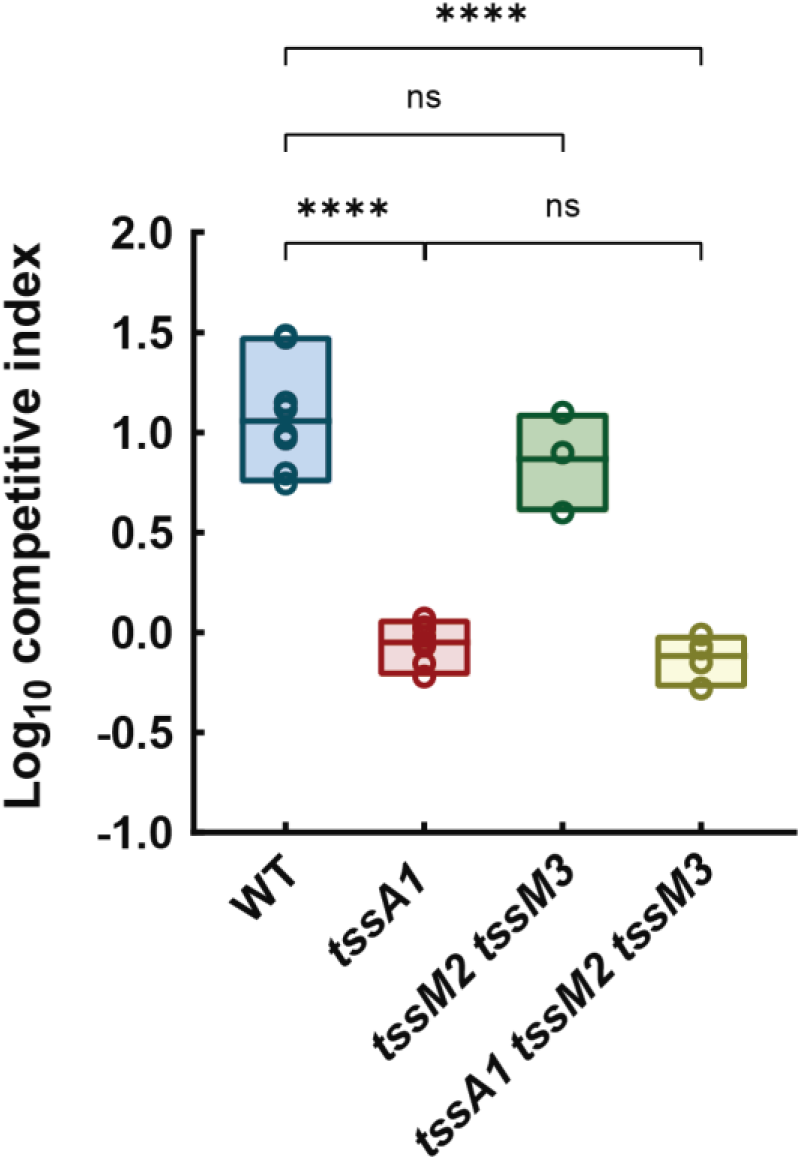
Competition assays between *P. putida* strains and *E. coli. P. putida* double mutant *tssM2 tssM3* does not show a decreased competitive index compared to the wildtype strain. Data are represented as floating bars showing values from the min to the max and with the line at the mean. Statistical analysis was performed using unpaired *t-tests.* N ≥ 3. P values are significant between the wildtype strain and the *tssA1* mutants.

### The K2/K3-T6SSs are used to kill plant pathogens

*P. putida* uses the K1-T6SS to eliminate ecologically relevant competitors, including severe plant pathogens such as *P. syringae*, *P. savastanoi* and *X. campestris* [17, 18]. We have observed that the K2- and K3-T6SSs are not active under standard laboratory conditions when *E. coli* is used as the prey in *in vitro* competition assays (Fig. 2). However, they are likely to be functional under other ecologically relevant conditions. To assess whether the K2- and K3-T6SSs are activated in the presence of competitors such as plant pathogens, we used the single, double and triple T6SS mutants described above to perform *in vitro* competition assays between KT2440 and its isogenic T6SS mutants and a representative plant pathogen, *X. campestris.* As previously reported, the *P. putida* wildtype strain caused a decrease in the survival of the tested phytopathogen with a competitive index of 4.5 [17, 18] and Fig. 3). As expected, the *tssA1* mutant has a significantly reduced competitive index when cocultured with the plant pathogen (Fig. 3) compared to the wildtype strain (C.I. ∼2.5). Interestingly, the same trend was observed in the double mutant *tssM2 tssM3* (C.I. ∼2.5) and the cumulative effect of the absence of all three systems was evident in the triple mutant (C.I. ∼1.5, Fig. 3). Unlike the previous observation of the competition assay against *E. coli,* where the K2- and K3-T6SSs remained silent, the competition assays between *P. putida* and *X. campestris* indicate that the K2- and/or K3-T6SSs are used to kill this phytopathogen. Growth experiments comparing mutant and wild-type strains demonstrated equivalent growth rate in the media used for competition assays (Fig. S3). This confirms that the observed differences between strains are not attributable to growth penalties or epistatic effects from the introduced mutations.

**Figure 3.**
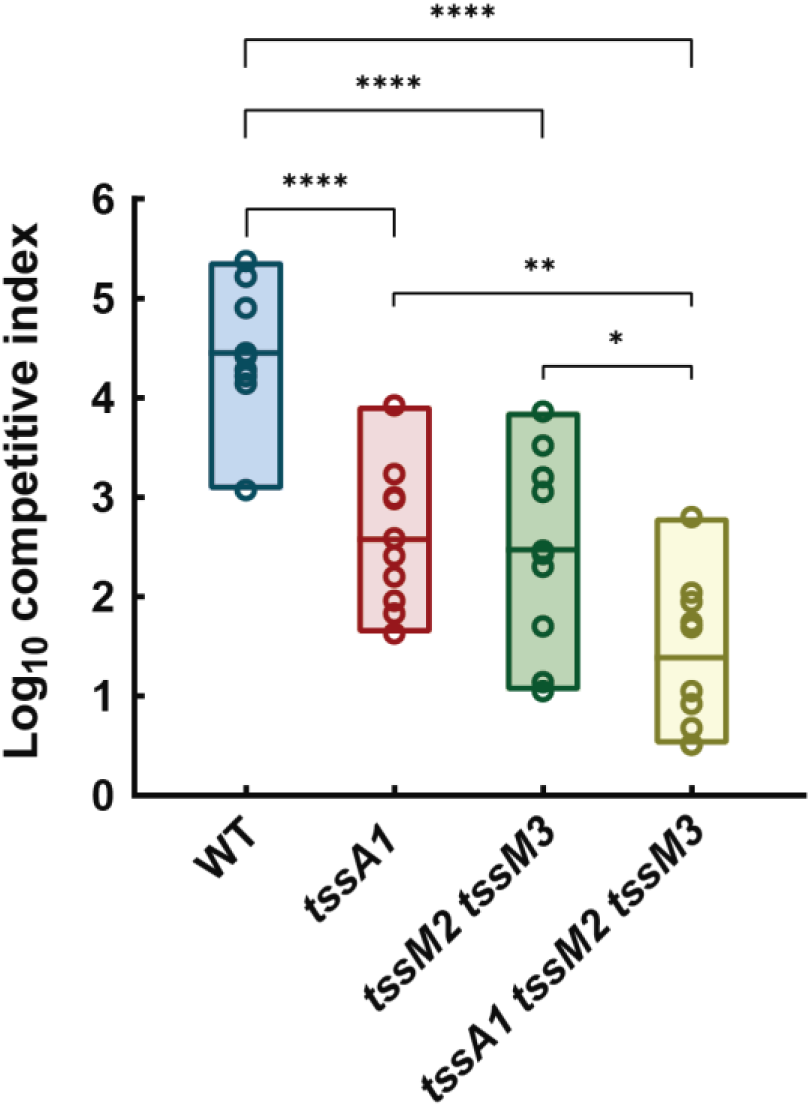
Competition assays between *P. putida* strains and *X. campestris. P. putida* double mutant *tssM2 tssM3* shows a decreased competitive index compared to the wildtype strain. Data are represented as floating bars showing values from the min to the max and with the line at the mean. Statistical analysis was performed using unpaired *t tests.* N ≥ 8. P values are significant between the wildtype strain and all the T6SS mutants.

### The *P. putida* T6SSs are key for rhizosphere colonisation

To determine the role of the T6SSs in the *P. putida* KT2440 colonisation of the rhizosphere, we inoculated 7-day-old tomato plants grown in microcosms containing non-sterile agricultural soil with *P. putida* and its T6SS mutant derivatives: *tssA1*, *tssM2 tssM3* double mutant and *tssA1 tssM2 tssM3* triple mutant. After a further 2-week incubation period, inoculated bacteria were isolated from the rhizosphere microcosms, by selecting with rifampicin. As shown in Fig. 4, inoculation with the mutant strains resulted in a significant reduction of 3-fold in bacterial recovery from the rhizosphere compared to the wildtype strain. This substantial decrease underscores the critical role that the K1-T6SS, K2-T6SS, and/or K3-T6SSs play in colonisation and persistence within the tomato rhizosphere microbiota. The diminished colonisation efficiency observed in the T6SS mutants suggests that these secretion systems are important for successful invasion and establishment within the rhizosphere. Colonisation of the rhizosphere by *P. putida* is particularly challenging due to competition from an established plant microbiota, making the T6SS a crucial mechanism that facilitates successful establishment and persistence in this competitive environment. The competitive advantage conferred by the T6SSs is likely decreased in the T6SS mutants because of their key role in interbacterial competition. By mediating competitive interactions, these systems enable *P. putida* to outcompete other microbial inhabitants in the rhizosphere environment.

**Figure 4.**
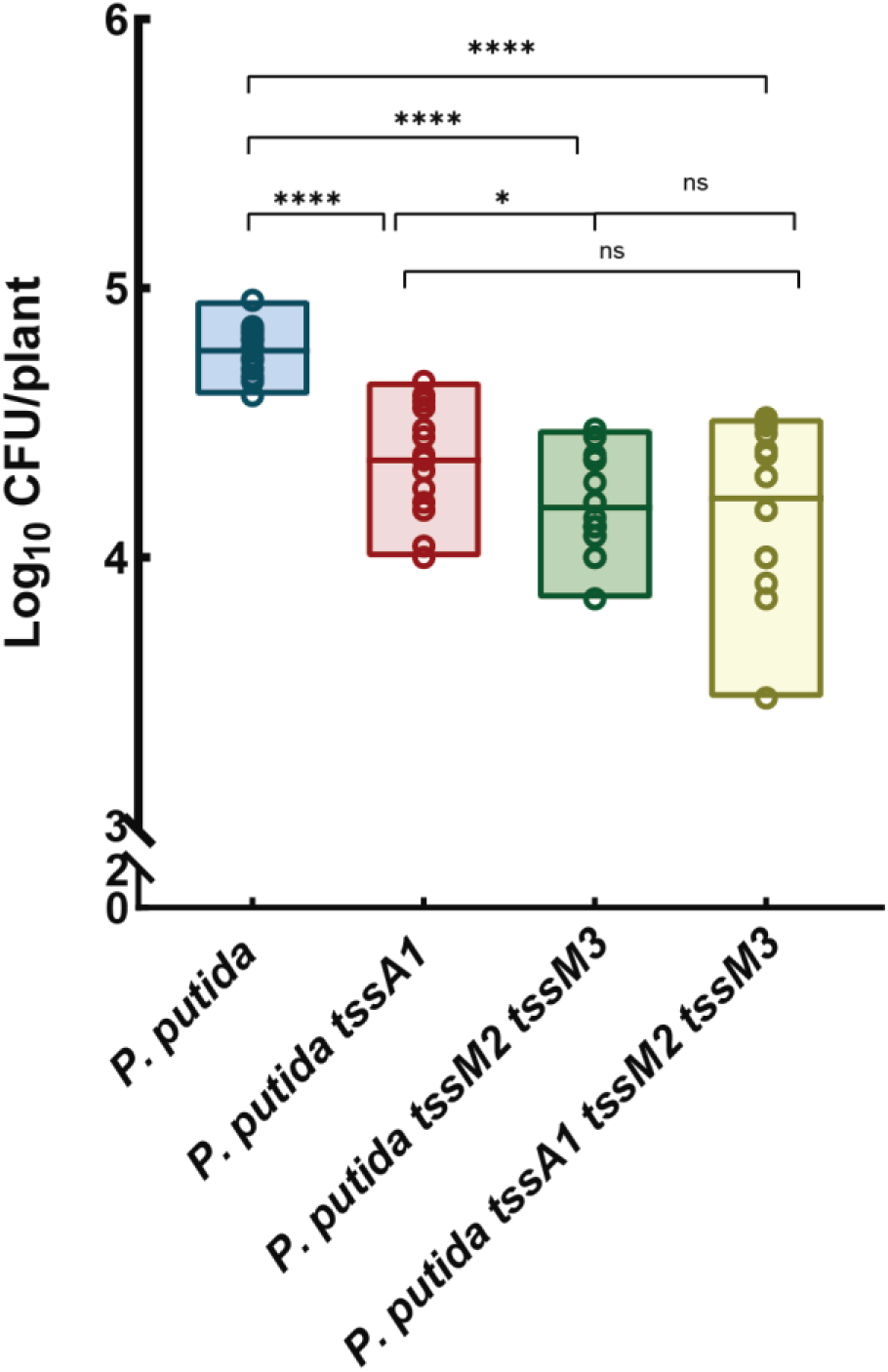
Quantification of bacterial recovery from rhizosphere colonisation. The figure shows the Log_10_ of the Colony Forming Units (CFUs) per plant recovered from the rhizosphere colonised by *P. putida* KT2440 and its isogenic mutant strains *tssA1*, *tssM2 tssM3* and *tssA1 tssM2 tssM3*. These strains were inoculated into the rhizosphere of tomato plants cultivated in non-sterile agricultural soil. Error bars represent the mean ± standard deviation (s.d.) of biological replicates. Statistical significance was determined using the Kruskal–Wallis non-parametric test (*P < 0.01, ** P < 0.001).

### The P. putida T6SSs shape the rhizosphere microbiota diversity

In a similar experiment, KT2440 and its isogenic mutants were separately inoculated into tomato plant rhizospheres. After a 2-week incubation period, DNA was extracted from the rhizosphere samples of the inoculated tomato plants and libraries of 16S DNA were constructed and custom sequenced.

To assess the impact of the T6SSs on the tomato plant microbiome, we calculated the Shannon index for each sample. This index is a common measure of alpha-diversity, reflecting both species richness (number of different species) and evenness (relative abundance of each species) within each sample. The analysis revealed no significant differences in the Shannon index among samples inoculated with the *P. putida* wildtype or the T6SS mutant derivative strains (Fig. 5A). These results indicate that the T6SSs do not influence the alpha-diversity of the microbial communities in the rhizosphere under these experimental conditions.

**Figure 5.**
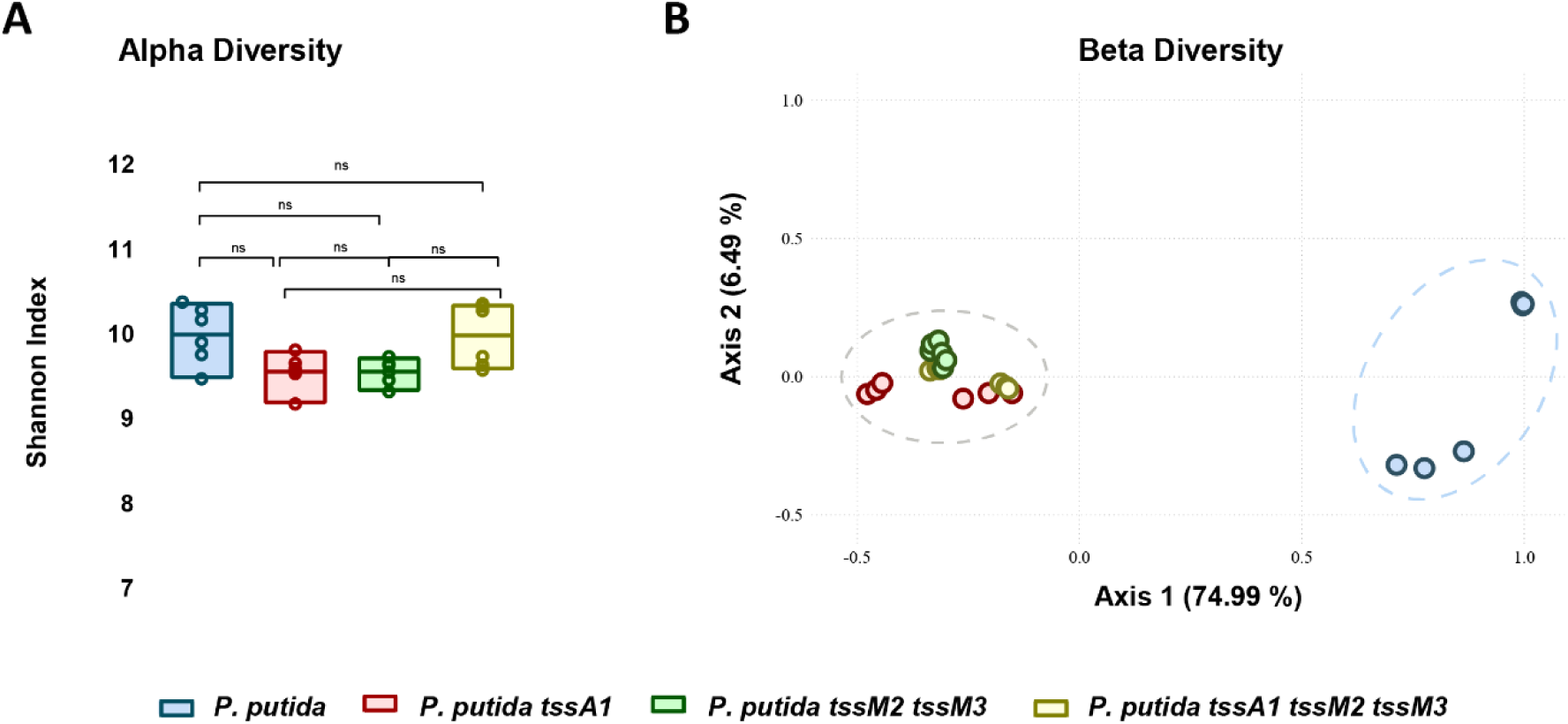
Analysis of rhizosphere microbiome diversity. **A.** Alpha-diversity was assessed using the Shannon index, and it shows no significant differences among inoculated samples. **B.** Principal Coordinates analysis (PCoA) of three 16S rRNA gene replicates from each soil sample using Bray-Curtis distances. Axis 1 explains 75 % of the observed variance.

However, the estimation of the beta-diversity by a principal coordinate analysis (PCoA) based on Bray-Curtis distances showed differences in the microbial communities’ composition between the treatments inoculated with the wildtype strain and the treatments inoculated with the T6SS mutants. The PCoA revealed that the first two Principal Coordinates explained a substantial portion of the variance in the data, with Principal Coordinate 1 accounting for 75 % and Principal Coordinate 2 for 6.5 %, together explaining 81.5 % of the total variance (Fig. 5B). The PCoA clearly shows a cluster of all samples inoculated with T6SS mutants separated from a second cluster containing those samples inoculated exclusively with the wildtype strain (Fig. 5B). Thus, the microbial community present in the rhizosphere inoculated with the wildtype strain is different from the communities found in the rhizosphere samples inoculated with the T6SS mutants, highlighting the role of the *P. putida* T6SSs in shaping polymicrobial communities such as the rhizosphere.

We performed a PERMANOVA (Permutational Multivariate Analysis of Variance) test to understand the influence of the T6SSs on the composition of the rhizosphere bacterial communities. Significant differences between the microbiomes of rhizospheres inoculated with the wildtype strain and those inoculated with the mutant strains were found (F= 23, *p-value* < 0.001, Table S4 and S5). Importantly, no significant differences were detected between the microbiomes of rhizosphere samples inoculated with the different mutant strains (Table S5). We also used ANCOM-BC to analyse the relative abundance of the bacterial phyla in the four samples (Fig. S4) and the differential abundance of specific bacterial groups at the family level (Fig. 6). The first analysis indicated that, among others, the abundance of the Actinomycetota phylum was higher in the rhizosphere samples inoculated with the T6SS mutants compared to the rhizosphere sample inoculated with *P. putida* wildtype strain (Fig. S4). A deeper analysis identified a wide range of taxa across multiple phyla significantly overrepresented in the rhizosphere microbiomes inoculated with the T6SS mutant strains compared to those inoculated with the wildtype strain (Fig. 6). As expected by the previous result, these taxa belong to the phylum Actinomycetota (Actinomycetota MB-A2-108, Cellulomonadaceae, Euzebyaceae, Gaiellaceae, Geodermatophilaceae, Ilumatobacteraceae, Intrasporangiaceae, Microbacteriaceae, Micrococcaceae, Micromonosporaceae, Nocardioidaceae, Solirubrobacteraceae, Solirubrobacterales bacterium 67-14, Streptomycetaceae, Streptosporangiaceae), but also to the phyla Acidobacteriota (Vicinamibacteria Subgroup 17, Vicinamibacteraceae), Bacteroidota (Chitinophagaceae), Chloroflexota (Anaerolineaceae, Chloroflexota Gitt-GS-136, Chloroflexota KD4-96), Cyanobacteriota (Nostocaceae), Entotheonellaeota (Entotheonellaceae), Bacillota (Aneurinibacillaceae, Bacillaceae, Paenibacillaceae), Myxococcota (Anaeromyxobacteraceae), Planctomycetota (Gemmataceae) and Pseudomonadota (Azospirillaceae, Beijerinckiaceae, Geminicoccaceae, Methyloligellaceae, Rhizobiaceae, Rhodobacteraceae) and represent the taxa impacted by the T6SSs of the wildtype strain. These findings suggest that the T6SSs of *P. putida* KT2440 play a crucial role in modulating the abundance and composition of diverse microbial groups in the rhizosphere.

**Figure 6.**
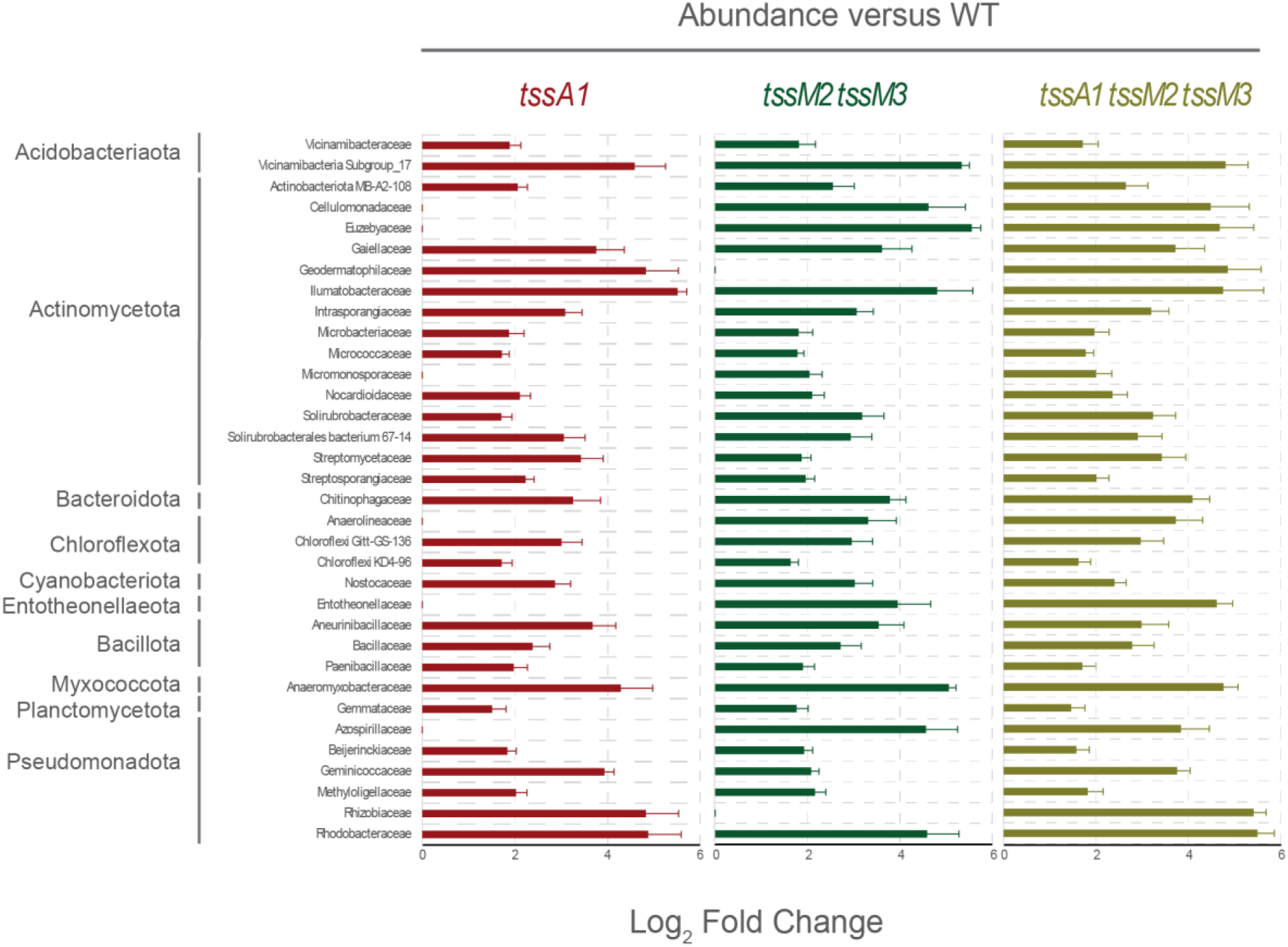
Differential Abundance Analysis of Rhizosphere Microbiome Taxa. The bar plot shows rhizosphere microbiome taxa significantly more abundant in the indicated T6SS mutants than in the wildtype strain, based on ANCOM-BC analysis (positive log_2_ fold change > 1.5, adjusted p < 0.001). The bar colours indicate the specific mutant strain compared to the wildtype: red (*tssA1*/WT), green (*tssM2 tssM3*/WT), and golden (*tssA1 tssM2 tssM3*/WT).

Importantly, our analysis reveals that the T6SSs of *P. putida* impact both Gram-positive and Gram-negative bacteria in the rhizosphere (Fig. 6). As expected, most of the phyla with taxa groups overrepresented in the T6SS mutant samples are predominantly Gram-negative phyla (Acidobacteriota, Bacteroidota, Chloroflexota, Cyanobacteriota, Entotheonellaeota, Myxococcota, Planctomycetota, and Pseudomonadota) however, other affected phyla such as Actinomycetota and Bacillota are Gram-positive (Fig. 6).

To investigate the putative specificity of the different *P. putida* T6SS clusters and their impact on rhizosphere microbiome composition, we employed a chord diagram based on differential abundance data derived from the ANCOM-BC analysis. The chord diagram allowed us to visualise the relationships between each T6SS group and the specific taxa they affect, providing information on how T6SS clusters contribute to shaping the composition of the microbial community by targeting specific bacterial groups (Fig. 7).

**Figure 7.**
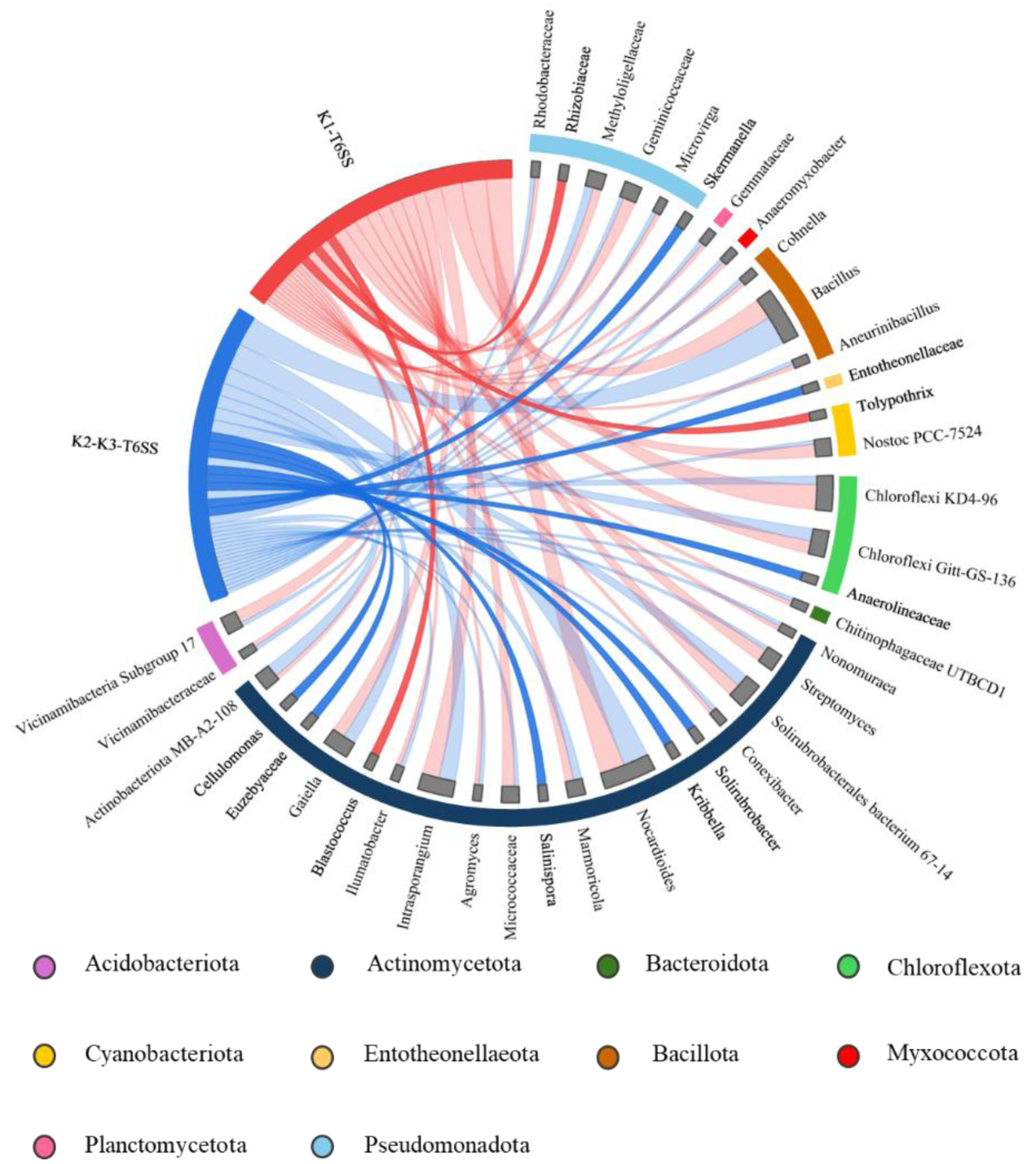
Taxa from the rhizosphere microbiome affected by the *P. putida* T6SSs and detected by ANCOM-BC analysis. **Grey bars**: all taxa affected by the T6SSs of *P. putida* KT2440. The length of the bars represents the number of unique ASVs detected for each taxon. **Red ribbon**: Taxa affected exclusively by the K1-T6SS. **Blue ribbon**: Taxa affected exclusively by the K2/K3-T6SSs. Three taxa are overrepresented in the samples inoculated with the strain lacking *tssA1* (*Rhizobiaceae*, *Tolypothrix* and *Blastococcus*). Eight other ASVs were overrepresented in samples inoculated with the strain lacking *tssM2* and *tssM3* (*Skermanella*, *Entotheonellaceae*, *Anaerolineaceae*, *Solirubrobacter*, *Kribella*, *Salinispora*, *Euzevyaceae* and *Cellulomonas*).

The analysis revealed that *Rhizobiaceae* and bacteria belonging to the genera *Tolypothrix* and *Blastococcus* are specifically targeted by the K1-T6SS, whereas bacteria from the genera *Skermanella*, *Entotheonellaceae*, *Anaerolineaceae*, *Solirubrobacter*, *Kribella*, *Salinispora*, *Euzevyaceae* and *Cellulomonas* are affected exclusively by the K2- and/or K3-T6SSs and not by the K1-T6SS. Most other taxa were similarly affected by the K1 and K2/K3-T6SSs.

## DISCUSSION

The T6SSs are present in 25% of all Gram-negative bacteria, being highly prevalent in the Pseudomonadota phylum (previously known as Proteobacteria) [6]. Many bacteria encode several phylogenetically distinct T6SSs, as is the case with *P. putida* KT2440, which encodes three systems [17]. Other pseudomonads encode from zero to three systems, with most strains encoding two or three [11, 12]. In other bacterial genera, such as *Burkholderia,* different species can harbour up to six distinct T6SSs [52], with four clusters conserved among three related species [53].

The presence of multiple phylogenetically distinct T6SSs within a single bacterial cell raises several questions regarding specialisation and spatial organisation. Firstly, understanding the degree of specialisation among these systems is crucial, *i.e.* whether each system possesses unique functions and/or targets or they randomly hit any close-by target. Secondly, the considerable size of a fully assembled T6SS poses a spatial challenge within the cell, leading to questions about how multiple systems can be accommodated and loaded simultaneously. While some T6SSs, such as the rapid *P. putida* K1-T6SS or *P. aeruginosa* H1-T6SS are assembled once per cell, others can coexist in multiple copies, with the slower *P. aeruginosa* H2-T6SS reaching more than four assembled systems per cell ready to be fired [18]. The activation of T6SSs is tightly regulated at transcriptional, translational and post-translational level [54] In *P. aeruginosa*, the phylogenetically distinct H1-, H2-, and H3-T6SSs are under the control of the same global regulatory network, GacA/GacS and Rsm with all three systems simultaneously present in an *rsmA* mutant [55]. Conversely, specific environmental cues can selectively trigger individual T6SSs, such as sub-optimal growth temperatures (25 °C) specifically inducing the *P. aeruginosa* H2-T6SS [56]. The *P. putida* K1-T6SS is activated during the stationary phase of growth or in the absence of global regulators RpoN, RpoS, or RetS [27], whereas the specific signals governing the K2- and K3-T6SS remained elusive until this study. Similarly, in *P. ogarae* F113, the F1- and F3-T6SSs are expressed in the rhizosphere and their expression is regulated by AmrZ and FleQ [12], while activation signals for the F2-T6SS are yet to be identified.

*P. putida* KT2440 contains three T6SSs (K1-, K2- and K3). The K2 and K3 share high sequence homology (Fig. 1) and belong to the same phylogenetic group [17]. However, the K2 system harbours mutations in *tssC2* and *vgrG2*, that introduce premature stop codons (Fig. 1), suggesting either inactivity or utilisation of paralogous proteins from the K3-T6SS.

Notably, the K2- and K3-T6SS clusters do not contain a *clpV* gene, which encodes the ATPase that recycles the sheath after the system fires. The ClpV component may not be necessary for these systems, as none of the clusters of the T6SS phylogenetic group 1.2a have a ClpV [11]. Functional T6SSs lacking *clpV* have been identified in other bacteria [57, 58] and *clpV* can be deleted without the total loss of T6SS function in *P. aeruginosa* or *V. cholerae* [59, 60].

Our study provides the first evidence that the K2- and/or K3-T6SSs, although they are not used to kill *E. coli* in *in vitro* competition assays, they are used to kill bacteria under ecologically relevant conditions. While the K1-T6SS is expressed in laboratory conditions and kills different Gram-negative bacteria, such as *E. coli* or plant pathogens [17, 18], the K2- and K3-T6SSs do not participate in *E. coli* killing (Fig. 2). However, they participate in *X. campestris* killing (Fig. 3). This demonstrates that the different T6SSs may be able to kill specific competitors, suggesting target specificity.

Similarly, the five T6SS clusters of *Burkholderia thailandensis* have been shown to exhibit distinct activities and target different competitors [53], though the differential impacts of these individual clusters on the host microbiome composition have not yet been investigated.

Previous research has shown that the T6SSs from *P. ogarae* and *P. chlororaphis* are involved in competition in the rhizosphere [12, 61]. Our results demonstrate that the *P. putida* T6SSs play a crucial role during tomato rhizosphere colonisation when other microorganisms inhabiting the natural soil are present (Fig. 4). The lack of the individual T6SS clusters results in a reduction of the abundance of *P. putida*. This reduction is significantly higher in a triple mutant with the three systems inactivated, indicating that the K1-T6SS and the K2- and/or K3-T6SSs are implicated in *P. putida* persistence in the rhizosphere environment.

Introducing *P. putida* in the rhizosphere system affects the resident microbiota (Fig. 5). While alpha-diversity is not significantly affected, beta-diversity analysis shows that the rhizospheric bacterial community influenced by the wildtype *P. putida* differs significantly from communities found in the presence of T6SS mutants. Notably, the absence of the K1-T6SS has a different impact on community structure than the absence of the K2/K3-T6SSs, suggesting system-specific effects on microbiome composition.

When *P. putida* colonises the tomato rhizosphere, it represents an introduced microbial community coalescing with the resident soil microbiota. The differential impacts of wildtype versus T6SS-deficient strains on community composition demonstrate how bacterial weaponry systems influence coalescence outcomes. The functionality of the previously dormant K2/K3-T6SSs specifically in response to plant pathogens and competitive pressure highlights an adaptive strategy that may determine success during community mixing events.

A wide range of taxa were significantly overrepresented in the rhizosphere microbiomes inoculated with the T6SS mutant strains compared with those inoculated with the wildtype strain (Figs. 6 and 7). This broad-spectrum influence suggests that *P. putida* uses its T6SSs to engage in competitive interactions across a wide phylogenetic range. Additionally, some taxa may be indirectly affected by the absence of T6SS activity through ecological cascades: while not directly targeted by *P. putida*, these taxa may benefit from reduced competitive pressure when their positively-interacting partners are no longer suppressed by T6SS-mediated killing, leading to their increased abundance in mutant-inoculated samples. The ability of the T6SSs to affect both Gram-positive and Gram-negative bacteria aligns with previous studies showing that T6SS effectors can include a variety of toxins capable of targeting both types of cell walls [62].

Our analysis reveals that the T6SS clusters of *P. putida* show specificity, with different taxa affected by the different T6SSs (Fig. 7). This indicates that the systems could target distinct bacterial populations, consistent with our observations for *E. coli* and *X. campestris* (Figs. 2 and 3). This dual mode of action (specificity and overlapping/complementary activity) suggests that *P. putida* employs a flexible strategy for competitive interactions within the rhizosphere, enhancing its adaptability and ecological fitness. These findings are consistent with Xiong et al., (2024), who demonstrated that the *P. putida* K1-T6SS drives compositional variation in multispecies biofilm communities [63].

In conclusion, we demonstrated that the *P. putida* T6SSs are crucial for its persistence in the tomato rhizosphere microbiota, with the first evidence of the K2/K3-T6SSs activation under ecologically relevant conditions. Wildtype *P. putida* significantly alters rhizosphere bacterial community structure compared to T6SS mutants, indicating the involvement of these systems in competitive interactions across diverse taxa. Our observation that different T6SS clusters may target specific bacterial populations highlights the flexibility of *P. putida* competitive strategy.

Further research exploring how the T6SS activity modulates community coalescence dynamics across different soil types and plant species would provide valuable insights into the fundamental ecological processes shaping rhizosphere microbiomes. Moreover, identifying the specific effectors deployed by each T6SS cluster and their respective bacterial targets would shed light on the molecular mechanisms underlying these microbiota-shaping interactions.

## ACKNOWLEDGEMENTS

M.M. acknowledges funding from the Spanish Ministry of Science, Innovation and Universities (MICIU/AEI/10.13039/501100011033) through the research grant from the State Subprogram for Knowledge Generation PID2021-125070OB-I00 (ERDF/EU). DV receives financial support from the Universidad Autónoma de Madrid through the PhD student fellowship for Research Personnel Training, FPI, program (FPI-UAM-SFPI/2021-00458).

P.B. acknowledges the financial support received from the Spanish Ministry of Science, Innovation and Universities (MICIU/AEI/10.13039/501100011033) through the Ramón y Cajal Program (RYC2019-026551-I, ESF Investing in your future), the research grant from the State Subprogram for Knowledge Generation PID2021-123000OB-I00 (ERDF/EU) and the research grant from the State Subprogram for Promotion of Research Consolidation CNS2022-135585 (European Union NextGenerationEU/PRTR).

We thank María Milagros López for her kind gift of the phytopathogenic strain.

## DATA AVAILABILITY STATEMENT

All data generated during this study that support the findings are included in the manuscript or the Supplementary data. Raw sequences fastq files are available in NCBI Bioproject PRJNA1064703.

## AUTHOR CONTRIBUTIONS (CRediT)

Conceptualisation: MM, RR, DD, PB

Data curation: DV, DD, PB

Formal analysis: DV, DD, PB

Funding acquisition: MM, RR, PB

Investigation: DV, DD, CC, AR

Methodology: DV, DD, PB

Project administration: MM, RR, PB

Resources: MM, RR, DD, PB

Software: DV, DD

Supervision: MM, RR, DD, PB

Visualisation: DV, PB

Writing—original draft: DV, PB

Writing—review & editing: DV, MM, RR, DD, PB

