## Supplementary Data for "The *Pseudomonas putida* Type VI Secretion Systems Shape the Tomato Rhizosphere Microbiota"

**This file includes:**

Figure S1 to S4

Tables S1 to S5

Supplementary references

**SUPPLEMENTARY INFORMATION FIGURES**

### Figure S1

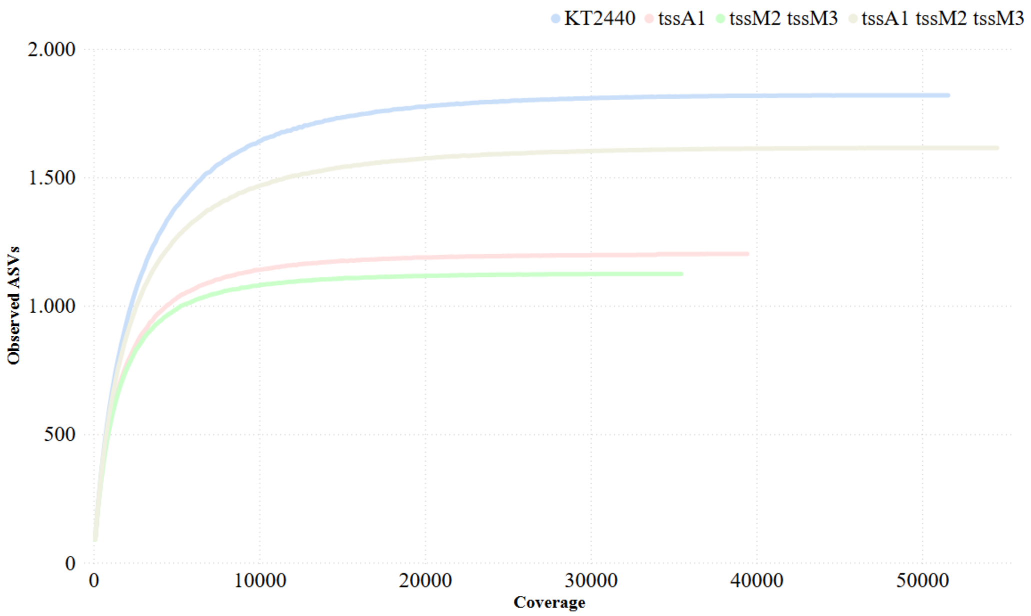


**Supplementary Figure 1 | Bacterial 16S rRNA Gene Amplicon Sequence Variant (ASV) Rarefaction Curves**. The plot represents the relationship between the number of sequenced reads (x-axis) and the corresponding observed diversity of bacterial ASVs (y-axis) for each of the 4 biological samples analysed. Each individual curve represents the combined sequence data obtained from the technical replicates of a single biological sample. All four curves plateaued, suggesting that the sequencing depth of our analysis was sufficient.

### Figure S2

**A)**

VgrG2-VgrG3-VgrG4-VgrG5 alignment

| Percent of Identity of VgrG2 with | **VgrG3** | VgrG4 | VgrG5 |
|---|---|---|---|
| | **96.31%** | 73.68% | 70.59% |

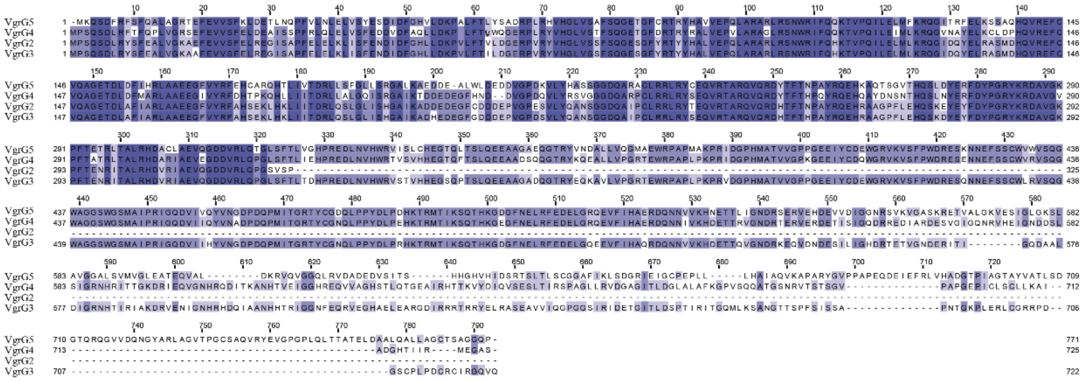

**B)**

TssC3-TssC2 alignment

Percent of Identity of TssC2 with TssC3 - **90.89%**

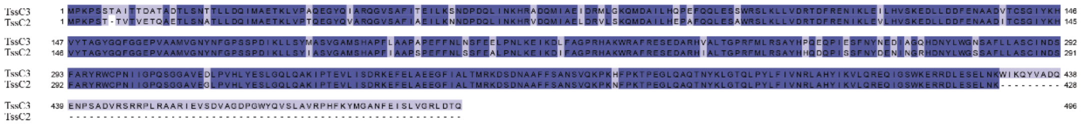

**Supplementary Figure 2 | Multiple Sequence Alignment (MSA) of** VgrG and TssC proteins of *P. putida* KT2440 performed with Clustal Omega (Maderia et al., 2022) and visualised by Jalview, coloured by percentage of identity. Simplified percent identity matrices (generated by Clustal 2.1) are included for each alignment (boxed in grey). **A)** VgrG proteins have a conserved N-terminal region (~1-550 aa), which allows interaction with other structural components of the system, primarily with the Hcp ring (Allsopp and Bernal 2023). The aligned VgrG proteins (VgrG2, VgrG3, VgrG4 and VgrG5) all contain the highly conserved region, but VgrG2 presents a premature stop codon that stops translation after amino acid 325. The identity between VgrG3 and the truncated VgrG2 is extremely high (>96%). **B)** Similarly, the identity between TssC3 and TssC2 is high (>90%) but TssC2 does not have the last 64 amino acids from a total of 495.

**Figure S3**

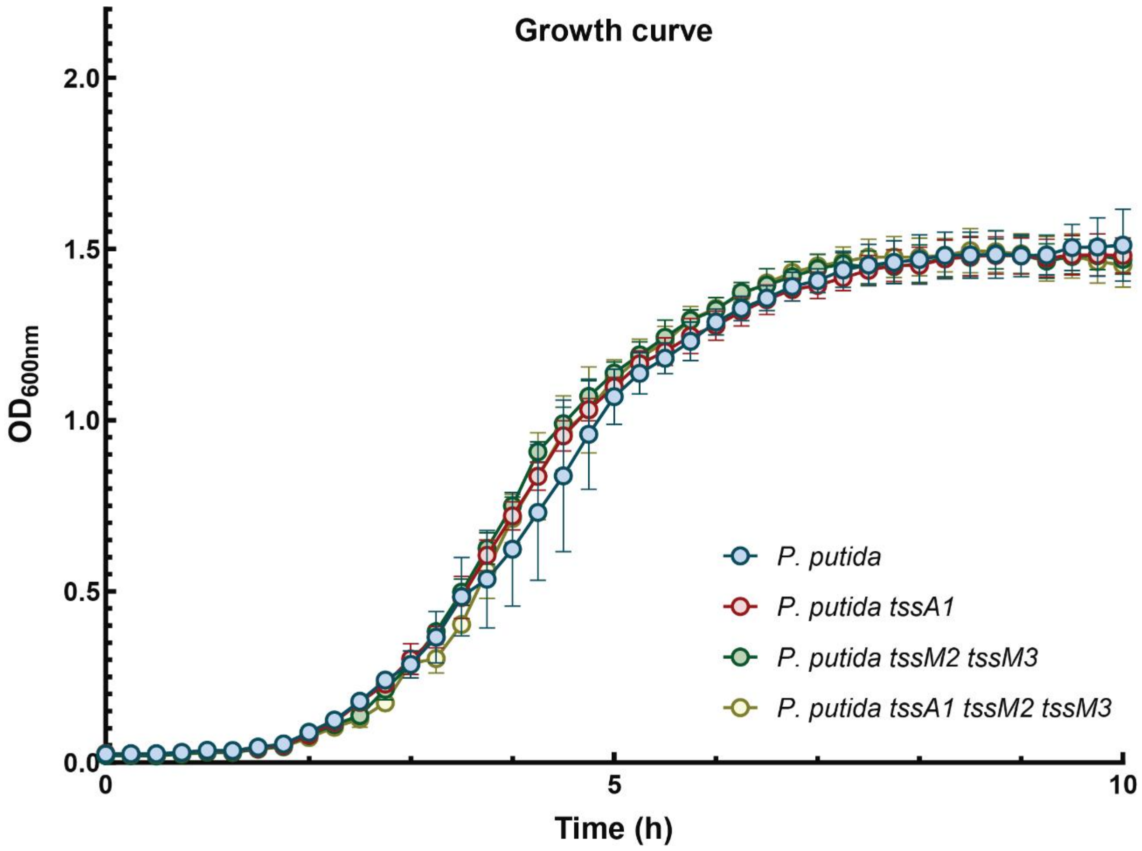


**Supplementary Figure 3 | Growth curve of *P. putida* wildtype and mutant strains.** Overnight cultures grown on LB Rif were adjusted to $OD_{600}$ of 0.05 in fresh LB media. $OD_{600}$ was measured for a total of 10 hours.

**Figure S4**

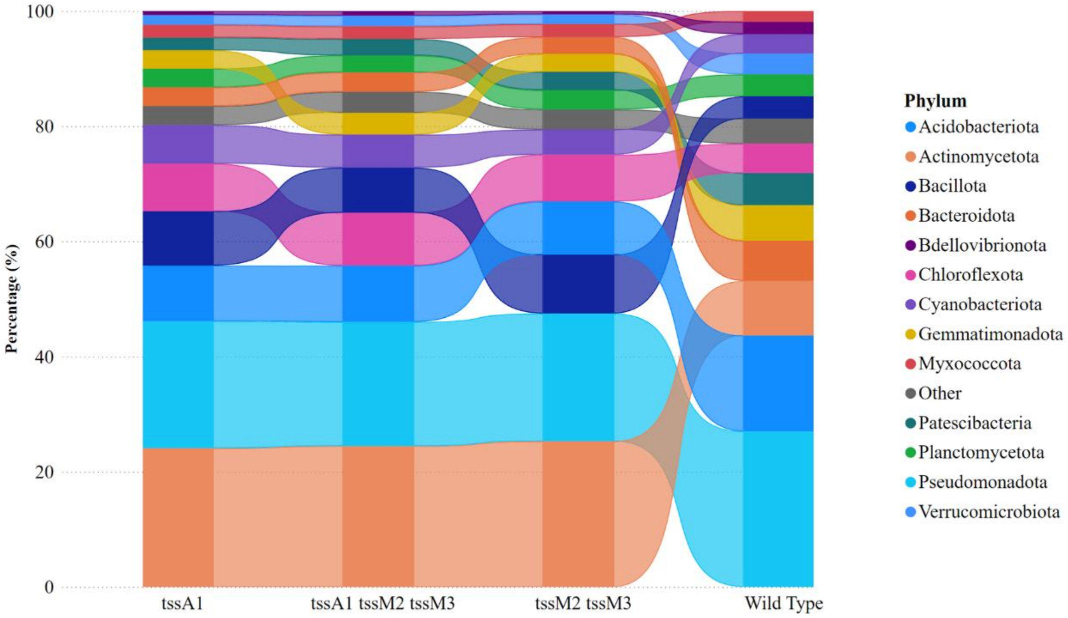


**Supplementary Figure 4 | Relative abundance of Bacterial Phyla in Inoculated Soil Sample**. The barplot represents the average relative abundance of bacterial ASVs aggregated at the phylum level, as observed in each inoculated soil sample two weeks post-inoculation. Each bar represents a specific soil sample inoculated with the wildtype, the *tssA1*, the *tssM2 tssM3* or the *tssA1 tssM2 tssM3* mutant strains. The height of the coloured segments within each bar indicates the average proportion of ASVs assigned to different bacterial phyla within that sample. To facilitate the comparison of individual taxa across the different samples, ribbons connect the segments representing the same phylum in adjacent bars.

### SUPPLEMENTARY INFORMATION TABLES

**Table S1.** Bacterial strains used in this study. The antibiotic resistance markers are identified as follows: Amp, ampicillin; Km, kanamycin; Gm, gentamicin, Sm, streptomycin, Nal, nalidixic acid; Pip, piperacillin and Rif, rifampicin.

| Name | Description | Source |
|---|---|---|
| ***Escherichia coli*** | | |
| DH5α | $F^-$ *end*A1 *gln*V44 *thi*-1 *rec*A1 *rel*A1 *gyr*A96 *deo*R *nup*G *pur*B20 φ80d*lacZ*ΔM15 Δ(*lac*ZYA-*arg*F*)*U169 *hsd*R17($r_K^- m_K^+$) $\lambda^-$, Nal$^R$ | [1] |
| CC118λpir | *ara*D Δ(*ara*, *leu*) Δ*lac*Z74 *pho*A20 *gal*K *thi*-1 *rsp*E *rpo*B *arg*E *rec*A1 λ*pir*, Rif$^R$ | [2] |
| HB101 | *supE44 hsdS20 recA13 ara-14 proA2 lacY1 galK2 rpsL20 xyl-5 mtl-1*, Sm$^R$ | [3] |
| ***Pseudomonas putida*** | | |
| KT2440R | Wild type strain, Rif$^R$ | [4] |
| KT2440R *tssA1* | Markerless mutant in the *tssA1* (PP3088) gene, disabling the K1-T6SS; Rif$^R$ | [5] |
| KT2440R *tssM2 tssM3* | Markerless KT2440R null mutant in the *tssM2* (PP4071) and *tssM3* (PP2627) gene, disabling the K2-and K3-T6SSs, respectively; Rif$^R$ | This study |
| KT2440R *tssA1 tssM2 tssM3* | Markerless KT2440R null mutant in the *tssA1*, *tssM2* and *tssM3* genes | [5] |
| ***Plant pathogens*** | | |
| *Xanthomonas campestris* pv. campestris IVIA 2734-1 | | [5] |

**Table S2.** Plasmids used in this study. The antibiotic resistance markers are identified as follows: Amp, ampicillin; Km, kanamycin; Gm, gentamicin, Sm, streptomycin, Cm, chloramphenicol, Pip, piperacillin and Rif, rifampicin.

| Name | Description | Source |
|---|---|---|
| pCR-BluntII-TOPO | Cloning vector, ColE1 *ori*, $Km^R$ | Invitrogen |
| pRK600 | Helper plasmid, ColE1 *ori, mob*RK2, *tra*RK2, $Cm^R$ | [6] |
| pKNG101 | Gene replacement suicide vector, R6K *ori*, *sacB;* $Sm^R$ | [7] |
| pKNG101-*tssM2* | 1.6-Kb PCR fragment containing the regions upstream and downstream *tssM2* cloned in pKNG101 (*Xba*I-*Bam*HI) for double recombination; when inserted into the chromosome and the plasmid cured, the strain is a *tssM2* mutant; $Sm^R$ | [5] |
| pKNG101-*tssM3* | 1.6-Kb PCR fragment containing the regions upstream and downstream *tssM3* cloned in pKNG101 (*Xba*I-*Bam*HI) for double recombination; when inserted into the chromosome and the plasmid cured, the strain is a *tssM3* mutant; $Sm^R$ | [5] |
| pRL662-*gfp2* | pRL662 [8] derivative expressing GFP2; $Gm^R$ | Erh-Min Lai collection |

**Table S3.** Oligonucleotide primers used in this study. The "Brief description" column provides basic information on the primer design (restriction enzyme used for cloning, encoded protein, forward or reverse orientation of the primer (F or R). Primers marked with UP or DOWN were used to construct mutant strains. Seq stands for sequencing primers. Primers tagged with 16S were used to amplify the V3-V4 regions of the 16S rRNA gene (**N**: any nucleotide; **W**: A or T; **H**: A or T or C; **V**: G or A or C). Sequences recognised by restriction enzymes are highlighted in lower case.

| Number | Brief description | Sequence (5´-3´) |
|---|---|---|
| P1 | XbaI.*tssM2*.UP.F | GAGAGtctagaGACGCGCCGAGCCATCTT |
| P2 | *tssM2*.UP.R | CTACATGACTTGATTCATCGAGGCTCC |
| P3 | *tssM2*.DOWN.F | ATGAATCAAGTCATGTAGGCAGGAGGC |
| P4 | BamHI.*tssM2*.DOWN.R | TTCAAggatccGCGTGAACGCTCGTTACA |
| P5 | *tssM2*.UP.F.Seq | TTGAGCTGGAGCGCCTGTTG |
| P6 | *tssM2*.UP.R.Seq | CGCGAGATCCGCTGGATAAC |
| P7 | XbaI.*tssM3*.UP.F | AGGAAtctagaACGACGCTACCGGCTACC |
| P8 | *tssM3*.UP.R | TCATAGTCGTTGATTCATCGAGGCTCC |
| P9 | *tssM3*.DOWN.F | ATGAATCAACGACTATGAACCTCGTCA |
| P10 | BamHI.*tssM3*.DOWN.R | ACCTTggatccTGAGCTGACGCTGCACAT |
| P11 | *tssM3*.UP.F.Seq | GAGCGCCTGCTAGGCAAGTAC |
| P12 | *tssM3*.UP.F.Seq | TGCGCGTCAACGGTATGTCG |
| P13 | TOPO.F.Seq | GTAAAACGACGGCCAG |
| P14 | TOPO.F.Seq | CAGGAAACAGCTATGAC |
| P15 | KNG.F.Seq | CATATCACAACGTGCGTGGA |
| P16 | KNG.R.Seq | CCCTGGATTTCACTGATGAG |
| P17 | 341F-16S | CCTACGGGNGGCWGCAG |
| P18 | 805R-16S | GACTACHVGGGTATCTAATCC |

**Table S4. Analysis of the influence of the *P. putida* T6SSs in the bacterial composition of tomato rhizosphere.** Assessment of the impact of T6SS mutations on the overall structure of the tomato rhizosphere microbiome performed by PERMANOVA (Permutational Multivariate Analysis of Variance) Adonis using distance matrices. The analysis partitions the total variation in the microbial community data to determine the proportion explained by the factor 'Strain' (wildtype and three T6SS mutants) versus unexplained residual variation. The results demonstrate a highly significant effect of strain (pseudo-F = 23.06, $p$ = 0.001), accounting for a substantial 77.6% of the total variation ($R^2$ = 0.776). This indicates a strong and non-random influence of the different strains on the multivariate composition of the rhizosphere microbiome. The table details the degrees of freedom (Df), sums of squares (SumsOfSqs), the pseudo-F statistic ($F$) value, the proportion of explained variance ($R^2$), and p-value derived from 9999 permutations (Pr(>F)), where a high F and low p-value signify significant differences attributed to the 'Strain' factor.

| PERMANOVA | **Df** | **SumsOfSqs** | ***F*** | ***R²*** | **Pr(>F)** |
|---|---|---|---|---|---|
| **Strain** | 3 | 0.223161 | 23.061843 | 0.775748 | 0.001 |
| **Residuals** | 20 | 0.064511 | NA | 0.224252 | NA |
| **Total** | 23 | 0.287671 | NA | 1.000000 | NA |

**Table S5. Pairwise PERMANOVA Analysis of Microbiome Differences Between Wildtype and T6SS Mutant Strains**. Following a significant global PERMANOVA test, pairwise comparisons were performed to pinpoint specific strain pairs with distinct microbiomes. The wildtype strain exhibited significant differences from all mutant strains, as indicated by high F and *p-values* < 0.005. In contrast, comparisons between the mutant strains were not statistically significant after multiple comparison correction, suggesting greater similarity among the microbiomes of these groups.

| Pairwise PERMANOVA | **Sample size** | **Permutations** | **F** | ***p-value*** |
|---|---|---|---|---|
| KT2440 vs *tssA1* | 12 | 9999 | 3.247927 | 0.0030 |
| KT2440 vs *tssM2M3* | 12 | 9999 | 4.148943 | 0.0023 |
| KT2440 vs *tssA1M2M3* | 12 | 9999 | 2.199285 | 0.0021 |
| *tssA1 vs tssM2M3* | 12 | 9999 | 1.149619 | 0.1046 |
| *tssA1 vs tssA1M2M3* | 12 | 9999 | 1.373882 | 0.0729 |
| *tssM2M3 vs tssA1M2M3* | 12 | 9999 | 1.050389 | 0.1604 |
